# A dynamic scale-mixture model of motion in natural scenes

**DOI:** 10.1101/2023.10.19.563101

**Authors:** Jared M. Salisbury, Stephanie E. Palmer

## Abstract

Some of the most important tasks of visual and motor systems involve estimating the motion of objects and tracking them over time. Such systems evolved to meet the behavioral needs of the organism in its natural environment, and may therefore be adapted to the statistics of motion it is likely to encounter. By tracking the movement of individual points in movies of natural scenes, we begin to identify common properties of natural motion across scenes. As expected, objects in natural scenes move in a persistent fashion, with velocity correlations lasting hundreds of milliseconds. More subtly, but crucially, we find that the observed velocity distributions are heavy-tailed and can be modeled as a Gaussian scale-mixture. Extending this model to the time domain leads to a dynamic scale-mixture model, consisting of a Gaussian process multiplied by a positive scalar quantity with its own independent dynamics. Dynamic scaling of velocity arises naturally as a consequence of changes in object distance from the observer, and may approximate the effects of changes in other parameters governing the motion in a given scene. This modeling and estimation framework has implications for the neurobiology of sensory and motor systems, which need to cope with these fluctuations in scale in order to represent motion efficiently and drive fast and accurate tracking behavior.

## INTRODUCTION

One of the great triumphs of theoretical neuroscience has been the success of the efficient coding hypothesis [1], which posits that sensory neural systems are adapted to the statistics of the organism’s natural environment. The importance of this hypothesis lies in its power to explain structural features of the nervous system, such as the shapes of nonlinear response functions [2] and receptive fields of sensory neurons [3–7], in terms of its function as an efficient information processing device. The success of this theory, particularly in vision, has sparked significant interest in measuring natural scene statistics (for a review, see [8]), and has continued to yield important results, like the ubiquity of non-Gaussian, heavy-tailed statistics and related nonlinear forms of dependency among scene features [9–12].

The observation of heavy-tailed distributions in the natural world connects with the rich structure that the external environment presents to an organism’s sensors, across a variety of sensory modalities. In any of these input streams, the brain has to pick out the relevant features in this rich input space that are most important for the organism’s survival–to select what matters. Adapting to this kind of structure and maintaining an efficient representation of behaviorally-relevant features in the world is a common feature of early sensory systems. Understanding how this is achieved, mechanistically, requires more than just the observation and quantification of heavy tails in natural scenes. To be able to understand how the brain represents this structure efficiently, we need to model it to shed light on potential ways the brain compresses this rich structure into an actionable internal signal.

Organisms are not passive sensory processors; they must also produce adaptive behavior in a complex and dynamic natural environment, where tasks like capturing prey [13–16], fleeing predators [17–19], and navigating obstacles [20] are all critical to survival. These behaviors inevitably involve prediction [21–24] in order to compensate for substantial sensory and motor delays [25]. The basis for such predictive behavior must be statistical regularities in the environment, but little is known about the statistics of the inputs relevant to such behaviors.

As a step towards characterizing the statistics of behaviorally relevant quantities in natural scenes, we focus on a feature fundamental to many essential sensation-to-action programs, the motion of objects. Object motion relative to the observer drives oculomotor tracking [26, 27] and is an essential part of many crucial behaviors, like prey capture [28–30]. Specialized circuitry as early as the retina distinguishes between object and background motion [31, 32], while entire brain regions in the visual cortex of primates specialize in processing motion [33], with increasing complexity along the dorsal stream [34].

While previous work has characterized motion in certain cases, often focusing on optical flow due to egomotion [20, 35, 36], little is known about the statistics of object motion in the natural world. To address this, we analyze movies from the Chicago Motion Database[37], which were shot and curated for the purposes of statistical analysis and for use as stimuli for neural recordings and visual psychophysics. Rather than trying to track discrete objects (which may be difficult even to define for some movies, like those of flowing water), we simplify the problem by tracking individual points within the image using classic techniques from computer vision [38, 39].

Given a point trajectory, the velocity along that trajectory is a spatially local description of an object’s motion through three-dimensional space, projected onto the two-dimensional surface of a sensor array, such as a retina or camera. We find that point velocity is highly correlated on the sub-second timescale we measure, and therefore point trajectories are highly predictable. More subtly, the distributions of velocity along trajectories exhibit heavy-tails and nonlinear dependencies, both across horizontal and vertical velocity components and across time.

This suggests the presence of an underlying *scale* variable, or local standard deviation, so the local velocity can be modeled as a Gaussian scale-mixture [40]. These models were developed in previous work examining the responses of filters applied to natural images and sounds [10, 41]. We find that the scale fluctuates within individual trajectories on a relatively short timescale, so it is an essential part of our description of natural motion. Despite considerable differences in the velocity statistics across movies, the dynamic scale-mixture structure is remarkably consistent. This has important implications both for the efficient encoding of motion signals by neurons–which must adapt to the fluctuating scale to make full use of their limited dynamic range [42–44]–and for behaviors relying on predictive tracking–which must take into account the highly non-Gaussian statistics of natural motion [45].

## RESULTS

In order to build up a statistical description of motion in natural scenes, we analyze movies from the Chicago Motion Database, which consists of a variety of movies collected for statistical analysis and for use as visual stimuli in experiments. All movies were recorded using a fixed camera, with scenes chosen to contain consistent, dense motion within the field of view for minutes at a time. Scenes include flowing water, plants moving in the wind, and groups of animals such as insects and fish. While natural visual input is dominated by the global optical flow during eye and head movements [20], object motion warrants specific attention because it is highly behaviorally relevant for essential behaviors like escape or prey capture. Note that these global and local motion signals are approximately additive, so one can combine them to form a more complete description of motion for a given organism. We analyze a total of 15 scenes, with a resolution of 512 × 512 pixels, each 2^14^ = 16, 384 frames long at a frame rate of 60 Hz (∼ 4.5 minutes). The high resolution, frame rate, and lack of compression of these movies are essential for getting precise motion estimates. We use lenses approximating the optics of animal eyes and provide rich metadata for each movie.

For each scene, we quantify local motion using a standard point tracking algorithm [38, 39]. A set of tracking points are seeded randomly on each frame, then tracked both forward and backward in time to generate trajectories (see *Materials and Methods* for details). Early visual and sensorimotor systems operate on a timescale of tens to hundreds of milliseconds, so we restrict our analysis to short trajectories (64 frames, or ∼ 1 s long) to reduce the amount of inevitable slippage from the point tracking algorithm. The resulting ensembles (2^13^ = 8, 192 trajectories each) sparsely cover most of the moving objects in each movie (Figure 1A).

**FIG. 1:**
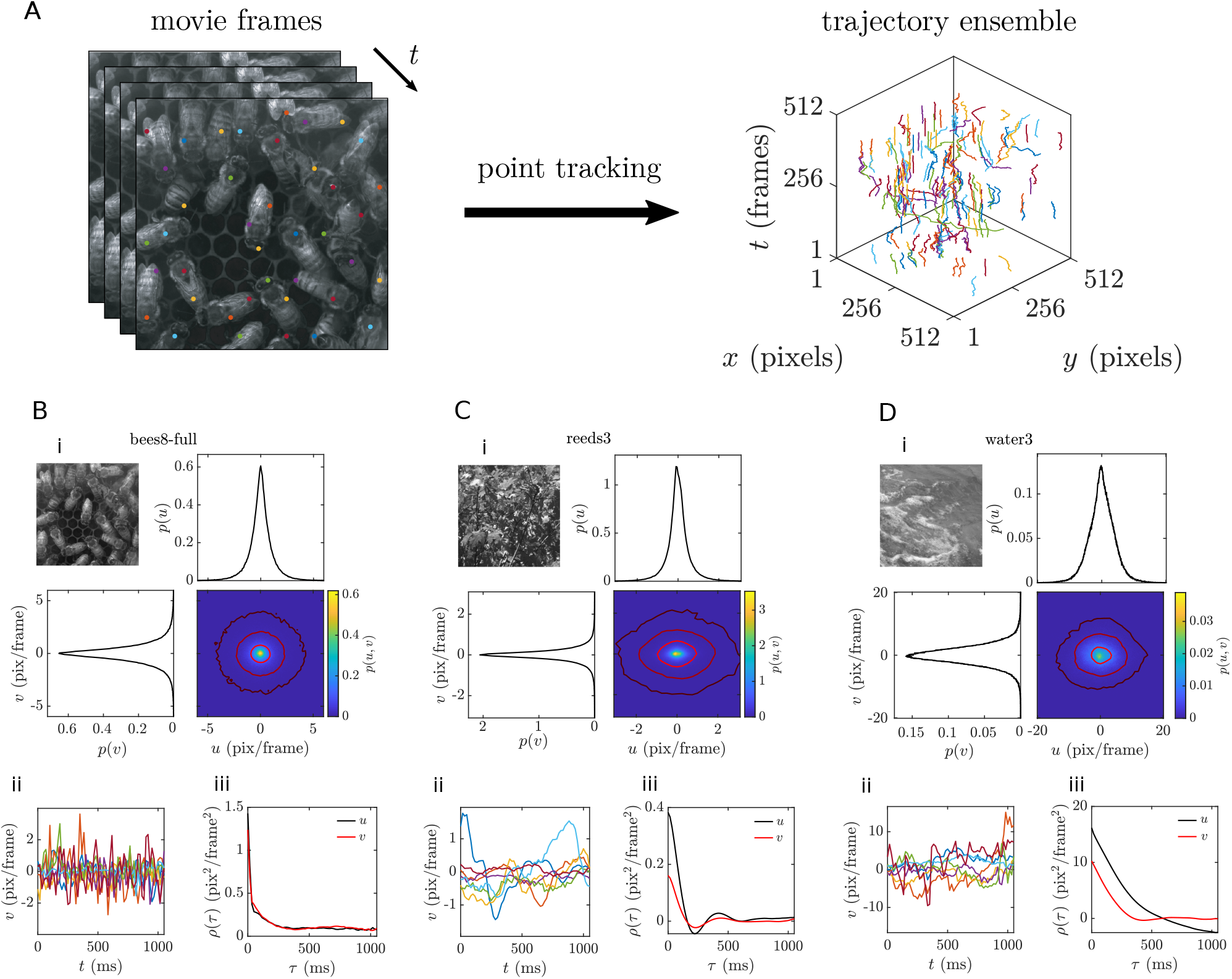
Automated point tracking reveals a diversity of motion statistics across natural scenes. **A**. Natural movie data analyzed via point tracking yields an ensemble of ∼ 1 s long point trajectories. **B-D**. Raw data summaries for three example movies, (**B**) `bees8-full`, (**C**) `trees14-1`, and (**D**) `water3`. **i**. Joint and marginal distributions for horizontal (*u*) and vertical (*v*) velocity components. Overlaid isoprobability contours for the joint distributions are *p*(*u, v*) = 10^−1^, 10^−2^, and 10^−3^ for **B** and **C** and *p*(*u, v*) = 10^−2^, 10^−3^, and 10^−4^ for **D. ii**. Seven example horizontal velocity component time series. **iii**. Horizontal and vertical velocity correlation functions.

The focus of our analysis is the point velocity, or difference in position between subsequent frames, measured in raw units of pixels/frame (this is easily converted to degrees of visual angle per unit of time, given a fixed viewing distance). The key advantage of this analysis is that the velocities are associated in time along a given point trajectory, which cannot be achieved by looking at the optical flow [46] or motion energy [47] alone. Note that since tracking is a difficult problem, the distribution of velocity constrained to good trajectories differs from the overall distribution, leading to underestimation of variance and kurtosis (see *Supporting Information*). This analysis is also distinct from previous work examining the spatiotemporal power spectra of natural scenes [48, 49], since power spectra measure the globally averaged pairwise correlations between pixels.

Our understanding of motion in natural scenes must be grounded in what is perhaps the first scientific study of motion in a natural setting: the diffusive motion of pollen particles in water observed by Brown [50], later described theoretically by Einstein [51] and Langevin [52]. See the *Supporting Information* for a discussion of Brownian motion and its relation to our modeling framework. Briefly, Brownian motion is characterized by a Gaussian velocity distribution with an exponential correlation function.

Natural scenes are, by definition, as richly varied as the natural world itself; each movie we analyze captures a small slice of this immense diversity. Our selection can be divided into three broad categories–animals, plants (animated by wind), and water–and we present summaries of the raw data for a representative movie from each category in Figure 1B-D. In contrast to the Gaussian velocity distributions expected for Brownian motion, histograms of the raw velocity data tend to be sharply peaked with long tails. Furthermore, the velocity time series exhibit correlation functions with diverse shapes, rather than a simple exponential decay.

### Heavy-tailed statistics of natural motion

To examine the heavy-tailed structure of the observed point-trajectory velocity distributions, we pool horizontal and vertical velocity components together for an example movie, bees8-full, and compare this histogram to a Gaussian distribution with the same variance (Figure 2A). Plotted on a log scale to highlight the tails, the empirical frequency falls off nearly linearly away from zero, while the Gaussian probability falls off quadratically. The same is true for the other movies in our dataset, pooled by category and all together (Figure 4A-C). Velocity distributions from animal and plant movies tend to have heavier tails, while those of water movies are closer to Gaussian.

**FIG. 2:**
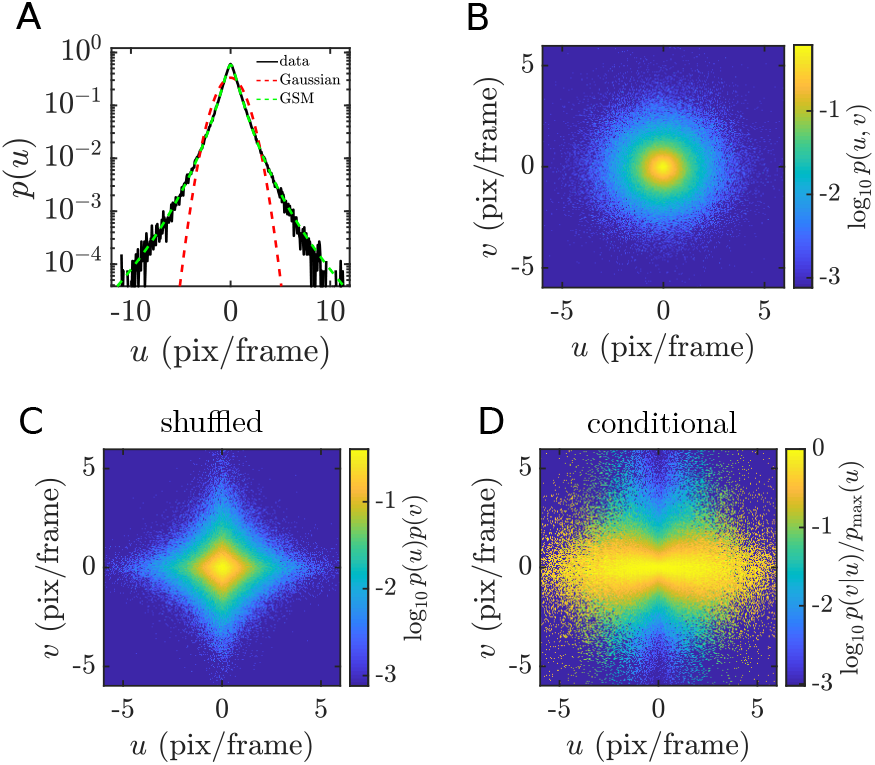
Velocity distributions are jointly heavy-tailed. **A-D**. Velocity distributions for a single example movie, `bees8-full`. The marginal distribution for horizontal velocity (**A**) has much heavier tails than a Gaussian with the same variance, and is well fit by a Gaussian scale-mixture model. The joint velocity distribution (**B**) is roughly radially symmetric, which differs substantially from the shuffled distribution (**C**) and indicates a nonlinear dependence between the two velocity components. This dependence is alternatively revealed by the conditional distribution of the vertical velocity given the horizontal velocity (**D**), showing a characteristic bow-tie shape.

When multiple variables are involved, heavy tails may be associated with a nonlinear form of dependency, as observed in the spatial structure of natural images [41]. The same is true for the two velocity components in our data. We illustrate this for `bees8-full`, but results are similar for all other movies. In Figure 2B we show a heat map of the joint histogram of horizontal and vertical velocity, *u* and *v*. It is nearly radially symmetric. (For other movies with unequal variance in the two components, distributions are elliptic.) When we shuffle th e da ta to break any association between *u* and *v*, the resulting histogram is no longer radially symmetric but is instead diamond-shaped (Figure 2C). This is a consequence of the fact that the Gaussian is the unique function which can be both radially symmetric and separable. We demonstrate this dependency more clearly by plotting the conditional distribution of *v* for each value of *u*, normalizing by the peak value at each *u* for visualization purposes (Figure 2D). The resulting “bow-tie” shape indicates that the variance of *v* conditioned on *u* increases with the magnitude of *u*.

The form of the velocity distributions observed above suggests that they can be modeled as Gaussian scale-mixture (GSM) distributions. As the name suggests, a GSM distribution is obtained by combining (zero-mean) Gaussian distributions of different scales, parameterized by a positive scalar random variable *S*. Let *Y* be a Gaussian random variable with mean zero and variance 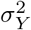. If *S* is a known quantity *s*, then *X* = *Y s* is simply a Gaussian random variable with mean zero and variance 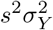. The conditional distribution is given by

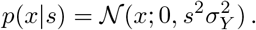

When *S* is unknown, *X* = *Y S* follows a GSM distribution given by

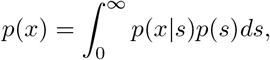

where *p*(*s*) is a distribution with positive support. A convenient choice is to let *S* = exp(*Z*), where *Z* is Gaussian random variable, which we will refer to as the scale generator, with mean zero and variance 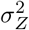. Then *S* follows a log-normal distribution, which simplifies the inference problem significantly, despite the fact that the resulting GSM distribution does not have a closed form. The choice of a log-normal distribution can also be justified by a maximum entropy argument [53]. See [41, 54] for a discussion of the GSM model in the context of wavelet analysis of natural images. Paremeters were estimated using a variant of the Expectation-Maximization (EM) algorithm [55] (see *Materials and Methods*).

For an individual velocity component as in Figure 2A, the GSM model captures the shape of the distribution well, with only two parameters: *σ*_*Y*_, controlling the over-all scale, and *σ*_*Z*_, controlling the heaviness of the tail. The variance of *X* is related to these parameters by

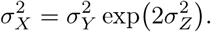

The kurtosis, which is the standard measurement of tail heaviness, depends only on *σ*_*Z*_:

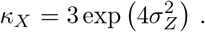

The kurtosis of *X* thus grows exponentially with the variance of *Z*, and matches the Gaussian kurtosis of 3 if and only if *σ*_*Z*_ = 0.

To model the joint distribution, as in Figure 2B, clearly we cannot use independent GSM models for each component, since this corresponds to the shuffled distribution in Figure 2C. Instead, we consider a model in which two independent Gaussian random variables, *Y*_1_ and *Y*_2_, are multiplied by a shared scale variable *S*:

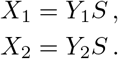

Note that we will maintain the general notation for the model for clarity. Applied to the velocity data, we have

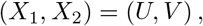

and (*Y*_1_, *Y*_2_) are the corresponding scale-normalized velocity components. The model is depicted in Figure 3, and it captures the radially symmetric (or more generally, when the variances are not equal, elliptic) shape of the joint distribution. This model has only three pa-rameters: 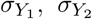, and *σ*_*Z*_ [56]. We observe a wide range of scale generator standard deviations *σ*_*Z*_ both within and across categories (Figure 4D). The trend across categories– namely that animal and plant movies tend to have higher scale standard deviations than water movies–agrees with the relative heaviness of the tails for the pooled data (Figure 4C). On the other hand, 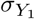 and 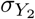 need not be similar, and many movies had a larger standard deviation of motion on the horizontal axis than vertical (the ratio 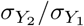 tended to be less than one, Figure 4E). The Akaike information criterion calculated for the two-dimensional GSM model with a common scale variable indicates that it is a better fit to the data than a two-dimensional, independent Gaussian model (Figure 4F).

**FIG. 3:**
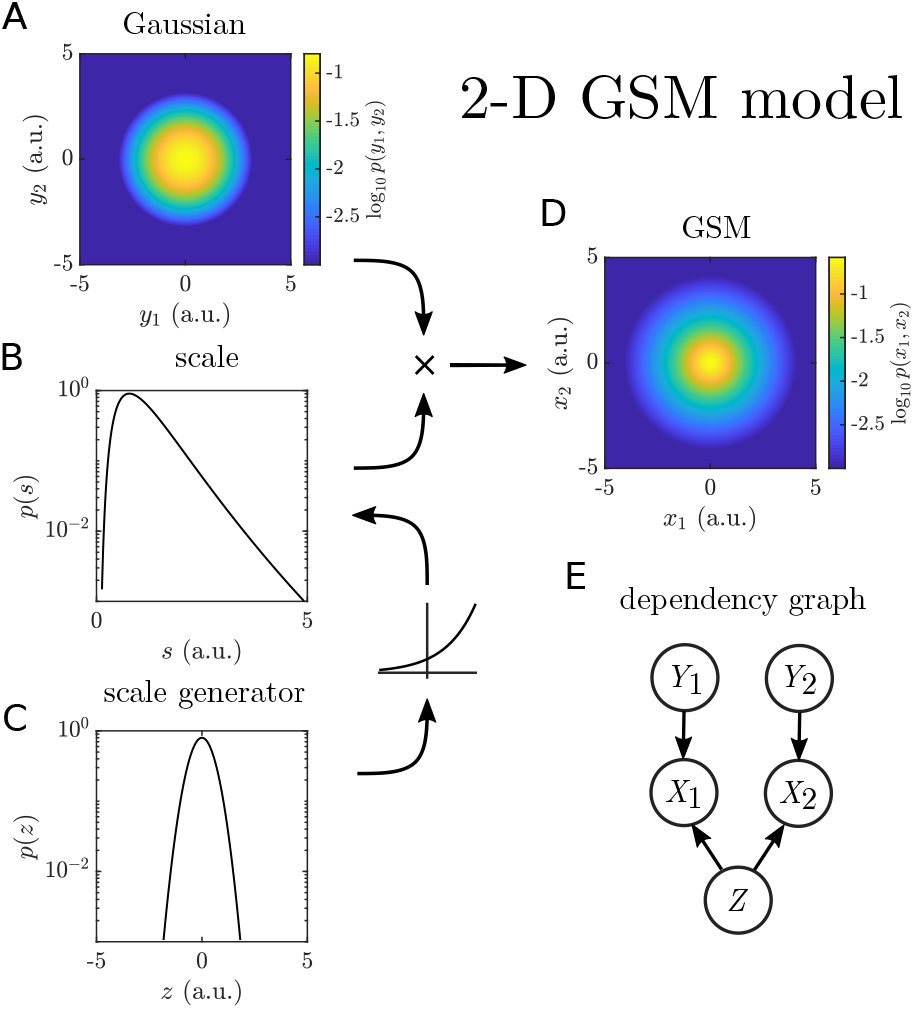
Schematic of the two-dimensional Gaussian scale-mixture model. **A-D**. A Gaussian random variable *Z* (**C**) is passed through an exponential nonlinearity to yield a log-normal scale variable *S* (**B**). The scale multiplies both components of an underlying Gaussian distribution (**A**) to produce radially symmetric heavy tails (**D**). For the joint distributions, probabilities less than 10^−3^ were set to zero to facilitate comparison with empirical histograms. **E**. Dependency graph for the variables in the model.

**FIG. 4:**
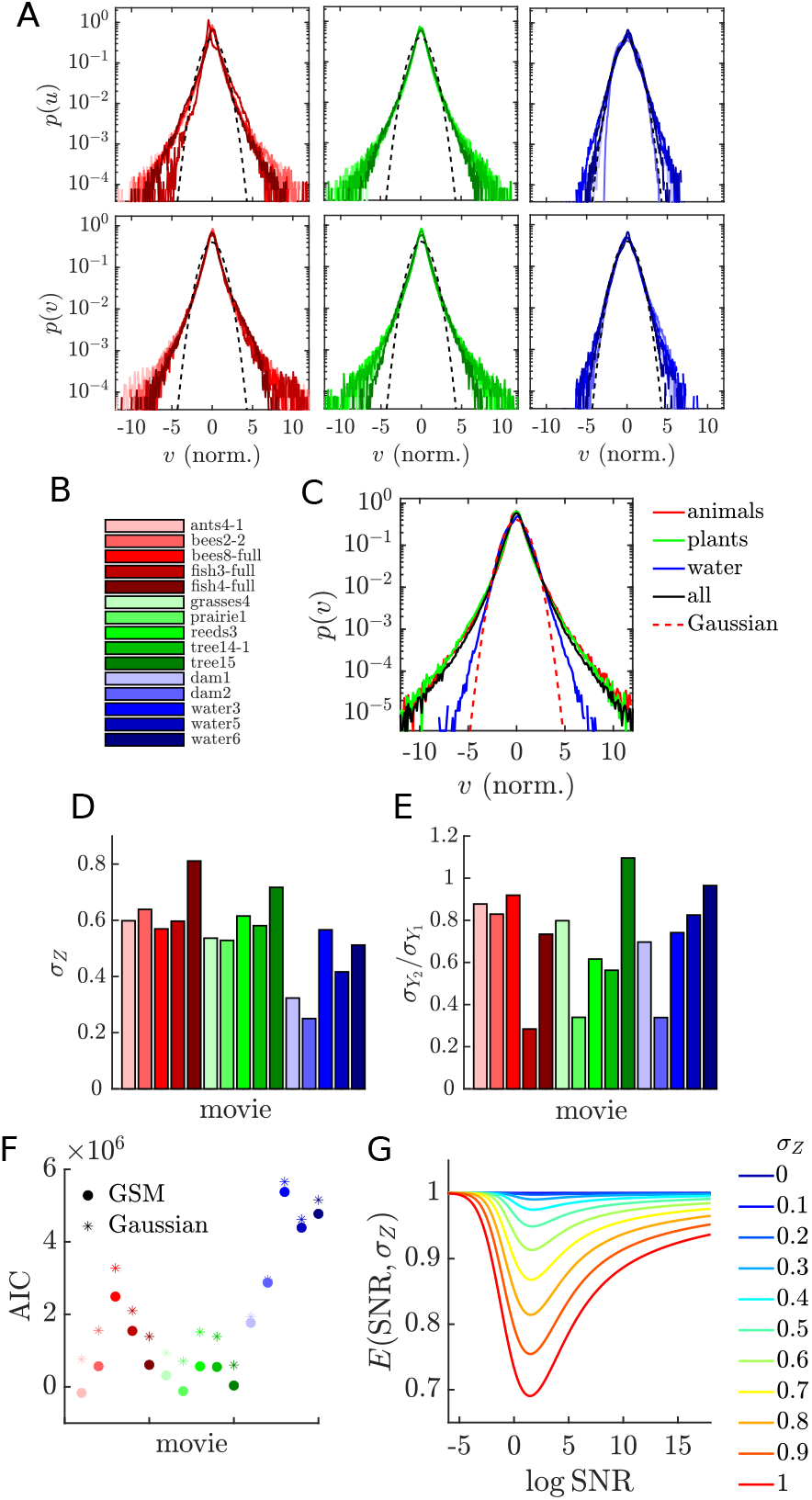
Quantifying heavy tails across scenes and categories. **A**. Marginal distributions for vertical and horizontal velocity components, grouped by category. **B**. Legend of individual movie names for **A** and all subsequent plots. **C**. Marginal distributions for the combined data across categories. Each velocity component of each movie was normalized by its standard deviation before combining. **D**. Estimated standard deviations for the scale generator variable, *Z*, varied across movies, corresponding to different amounts of kurtosis. **E**. The ratio of estimated standard deviations of the underlying Gaussian variables, *Y*_1_ and *Y*_2_, showing the the degree of anisotropy. **F**. AIC values for the two-dimensional, shared scale GSM model versus the two-dimensional, independent Gaussian model. **G**. Coding efficiency as a function of signal-to-noise ratio for different values of *σ*_*Z*_.

### Coding implications of heavy tails

Heavy-tailed velocity distributions pose a particular challenge for efficient coding via sensory neurons. Consider the classic information theoretic problem of coding a random variable *X* with an additive white Gaussian noise (AWGN) channel [57] [58]. The channel capacity, *C*, is a function of the signal-to-noise ratio

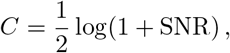

where 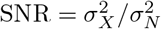 is the ratio of the signal variance to the noise variance. The mutual information, *I*, between *X* and its decoded estimate 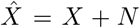, for Gaussian noise *N*, is equal to *C* if and only if *X* is Gaussian. Otherwise, *I* < *C*, and the coding efficiency *E* = *I/C* is less than one.

We calculate the coding efficiency given the parameters of a GSM model for *X* and the noise level 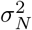 to explore the effects of heavy tails. We have

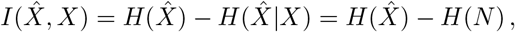

where

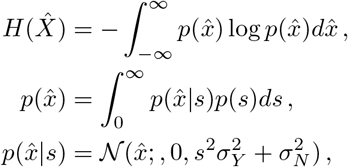

and

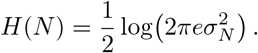

To see the effect of heavy tails, we vary 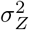 and SNR, keeping either 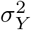 or 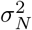 fixed (Figure 4F). Since *I* and *C* have the same scaling behavior with SNR, *E* → 1 as *SNR* → *±*∞, so the heavy tails have no effect at very high or low SNR. At intermediate SNR, the coding efficiency decreases monotonically as 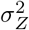 increases. The efficiency reaches a minimum at log SNR = 3*/*2 for all 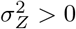.

The above calculation describes the loss of coding efficiency when *X* is sent through the channel with a constant gain. There are several ways to manipulate *X* for better efficiency. One is to ‘Gaussianize’ *X*, that is, to apply a compressive nonlinearity *f* such that *f* (*X*) is Gaussian. Neurons have been shown to implement this kind of efficient coding by matching their response nonlinearities to natural scene statistics [2], although the mapping is to a uniform distribution over a fixed interval rather than a Gaussian[59]. This method can be applied to each channel (velocity component) and time step independently. The downside of this strategy is that it introduces signal-dependent noise, since 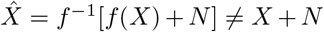. In particular, velocity values with high magnitude, which may be the most relevant for behavior, will have high noise.

Another strategy is to demodulate or normalize *X* by estimating *S* and dividing *X* by it. If the estimate is accurate, then 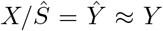, a Gaussian, and channel efficiency for 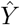 will be high. In order to recover *X*, 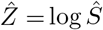 will need to be encoded in another channel, and the two sources of additive noise will result in multiplicative noise and heavy-tailed additive noise in the estimate:

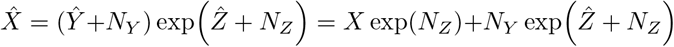

The question of whether it is better to use a single channel inefficiently or to use two channels efficiently depends on the cost associated with each channel and its SNR-dependent energy consumption.

Of course, this strategy fails for a single variable *X* since the only reasonable estimate for the scale is *Ŝ* = |*X*|, so that 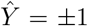. However, since two velocity components share a common scale variable, the estimate can be improved by making use of both components. Furthermore, since the scale is correlated in time, as shown in the next section, the history of *X*(*t*) can also be used to further improve the estimate *Ŝ*.

### The dynamics of natural motion

While the time-independent statistics of velocity are important, a full description of how objects move must include how the velocity evolves over time *along* point trajectories[60]. This motivates our point tracking analysis, which provides information that cannot be gleaned from motion estimates at fixed locations alone. From the raw data we know the velocity is highly correlated at short time lags, but it is not clear how the scale variable enters into play. We again inspect the joint velocity distribution for an example movie, now across neighboring time points for one velocity component (Figure 5A). The tilt indicates strong linear correlation across time in the velocity, as expected, and we note that the overall shape is elliptic, as in the uncorrelated GSM model. In Figure 5B we condition on the velocity at one time-step, and observe the same bow-tie shape as in the horizontal-vertical joint distribution. Thus, two forms of dependence–linear correlation and the nonlinear dependence due to the scale variable–coexist.

**FIG. 5:**
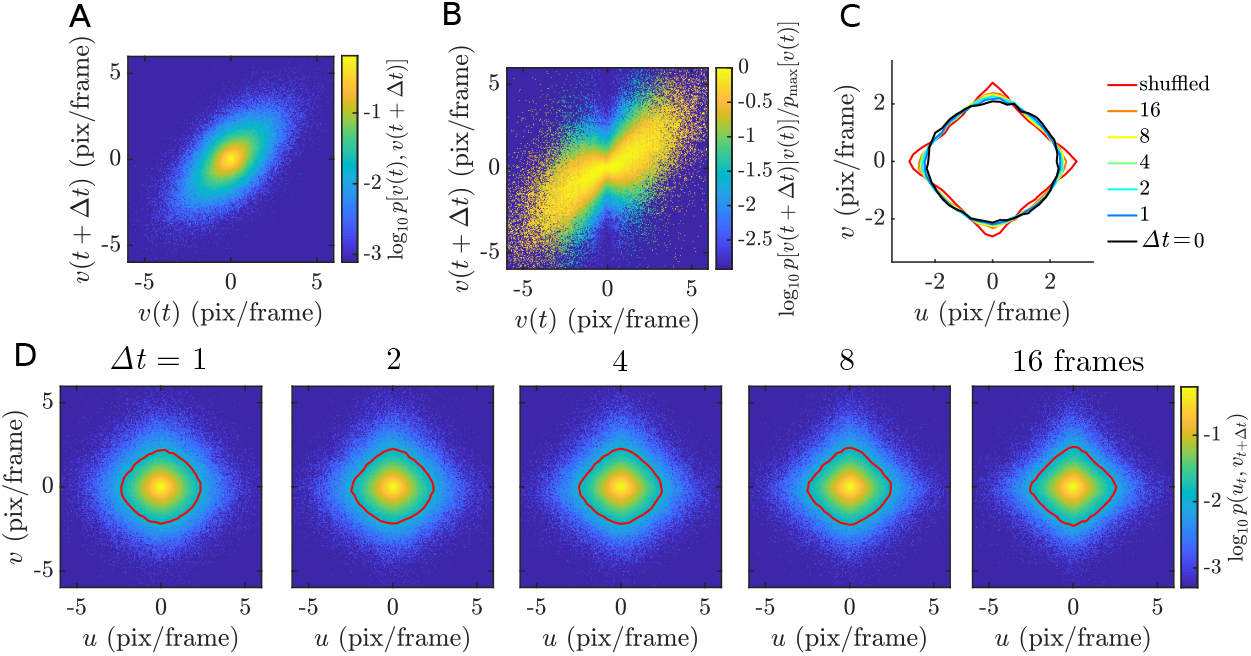
Temporal correlations in velocity and scale. **A-B**. Joint (**A**) and conditional (**B**) histograms for horizontal velocity across two adjacent frames for an example movie (bees8-full). The tilt indicates a strong linear correlation, while the elliptic shape in (**A**) and bow-tie shape in (**B**) indicate the coexistence of a nonlinear dependence due to an underlying scale variable. **C-D**. Isoprobability contours at *p* = 0.01 of the joint distributions of the two components separated by *τ* frames (**D**) show a gradual transformation from the original circle (**Figure 2B**) towards the diamond shape of the shuffled distribution (**Figure 2C**), indicating that the nonlinear dependence decays slowly over time. Isoprobability contours are overlaid in **C** for clarity.

We next ask whether this scale variable is constant in time (varying only from trajectory to trajectory) or dynamic (varying in time within a given trajectory). If it is constant, the jointly heavy-tailed distribution of the two components will not depend on the alignment of the two components in time, so long as they are from the same trajectory. In Figure 5D, we examine these joint distributions after shifting one component relative to the other by a time lag, for a range of lags. The distributions gradually shift from the radially symmetric zero-lag distribution to a diamond shape similar to the shuffled distribution. This is most clearly seen by comparing the *p* = 0.01 isoprobability contours as the lag increases (Figure 5C). In other words, the nonlinear dependence induced by the shared scale variable decreases as the lag increases. We conclude that the underlying scale variable is dynamic. Notably, we can detect these changes within the ∼ 1 s long trajectories to which we limit our analysis. This would not be the case if the scale were to change only on a very long timescale or only across different point trajectories within a scene.

We would like to capture this dynamic scale variable in our model of natural motion. It is straightforward to make the GSM model dynamic by replacing each Gaussian variable with an autoregressive Gaussian process, and we call this new model the ARGSM model. We illustrate it schematically in Figure 6 by generating exam-ple traces for one Gaussian velocity component *Y* and the scale generator *Z*. Note that the autoregressive scale generator variable is the temporal equivalent to the spatial Markov random fields explored in the image domain [41, 61, 62]. Given this model, we perform estimation of the parameters using a stochastic approximation variant of the expectation-maximization (EM) algorithm (see *Materials and Methods*). Example traces illustrating the results of this model are shown in Figure 7A-C. The estimated autoregression coefficients determine the correlation functions of the underlying Gaussian velocity and scale generator processes, which we plot for each movie in Figures 7D and E, respectively. The fact that some velocity correlation functions and many scale generator correlation functions do not go to zero over length of the trajectories could indicate a nonzero mean component that varies from trajectory to trajectory, but this is beyond the scope of the present analysis. Average correlation functions across categories are shown in Figure 7F-G. We also report the time to 0.5 correlation for each movie for the velocity in Figure 7H-I. Akaike information criterion (AIC) scores (Figure 7K) indicate that the full ARGSM is a better fit to the data compared to the AR model. It is also a better fit compared to the ARGSM model with a static *Z* value for each trajectory, indicating that a dynamic scale variable is essential for describing the data. In the context of visual tracking of moving objects, the timescales of these correlations functions are extremely important. On one hand, the velocity correlation time determines how far into the future motion can be extrapolated. On the other hand, the scale correlation time determines the timescale on which adaptation must take place in order to efficiently process motion signals with limited dynamic range.

**FIG. 6:**
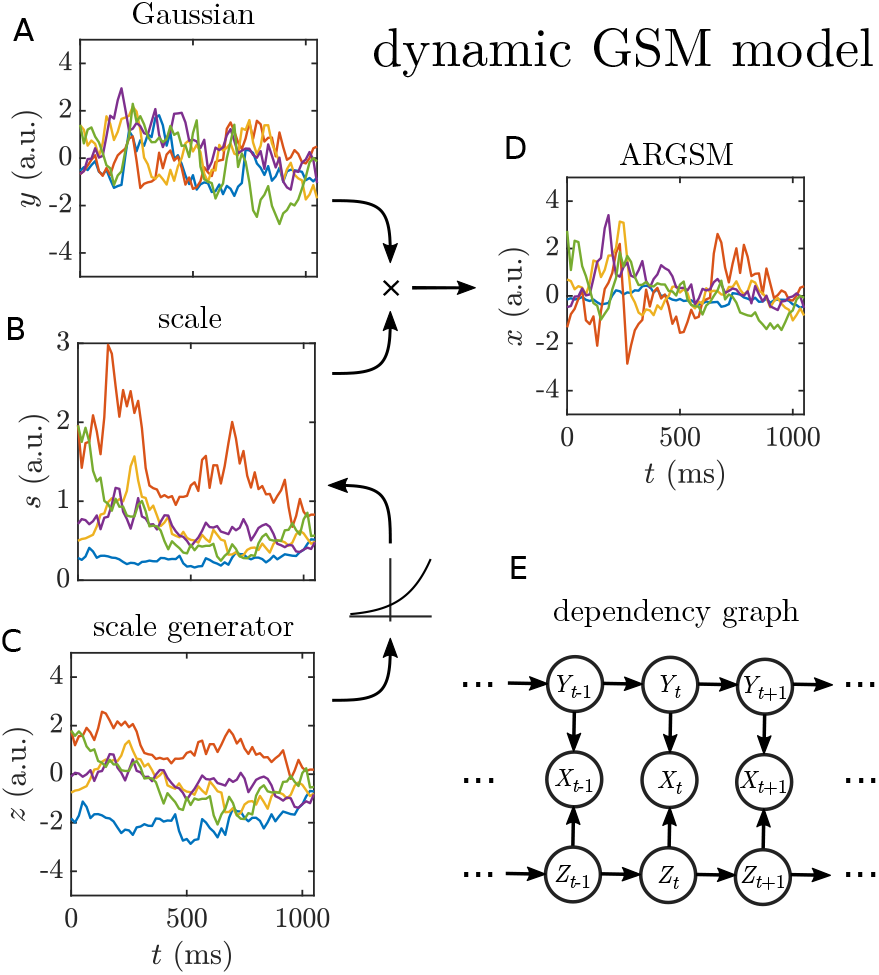
Schematic of the dynamic Gaussian scale-mixture model. **A-D**. Both *Y* (**A**) and *Z* (**C**) are modeled by high-order autoregressive processes to capture arbitrary correlation functions. (Only AR(1) processes are depicted graphically and used to simulate data.) The scale process *S* (**B**) is generated by passing *Z* through an element-wise exponential nonlinearity. It then multiplies the underlying Gaussian process *Y* element-wise to yield the observed process with fluctuating scale (**D**). Only one component is depicted. In the full model, two independent Gaussian processes share a common scale process. **E**. Dependency graph for the variables in the one-dimensional model.

**FIG. 7:**
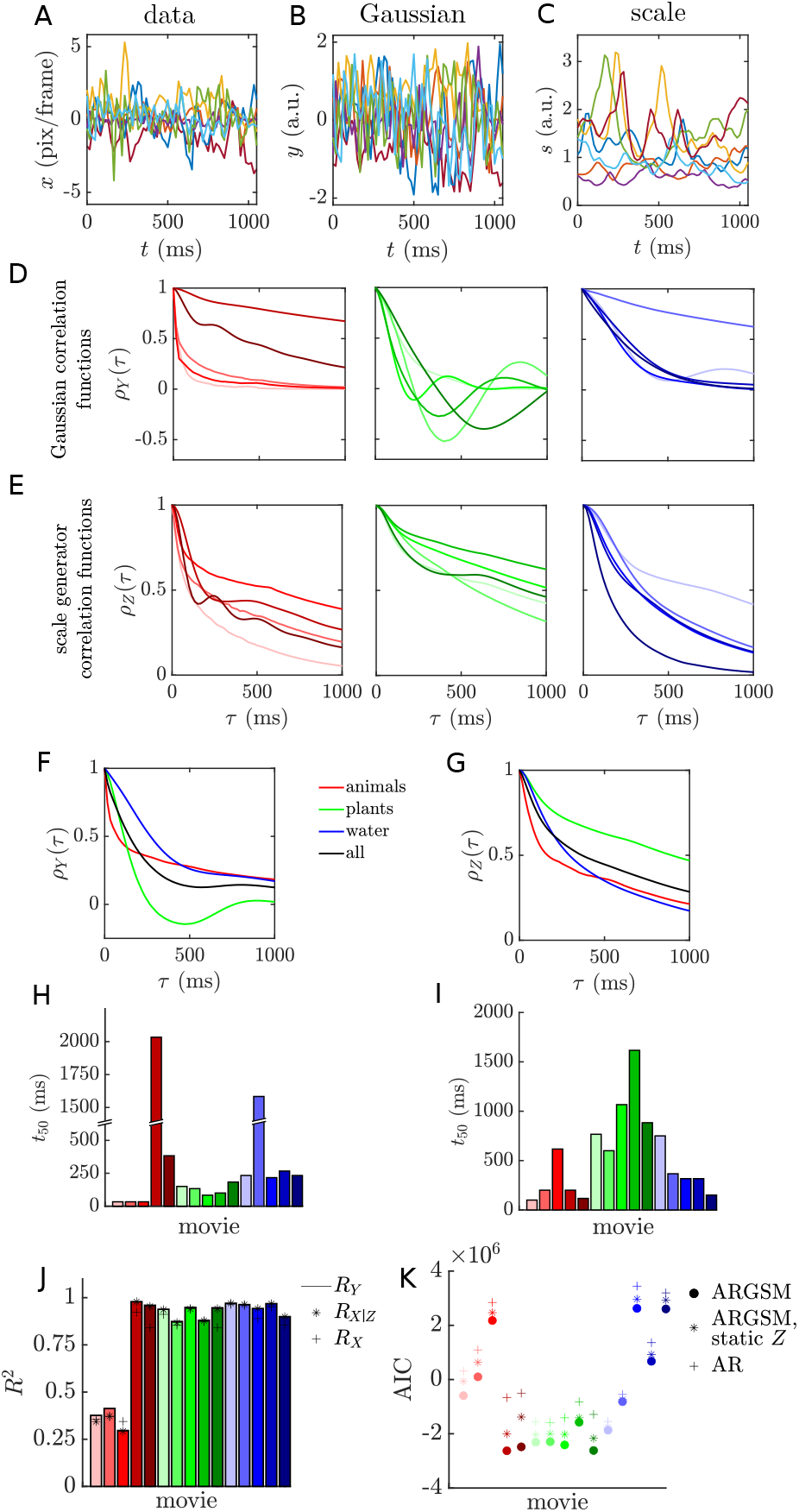
Quantifying velocity and scale correlations. **A-C**. Example traces of the raw velocity (**A**), scale-normalized velocity (**B**), and estimated scale variable (**C**). **D**. Temporal correlation functions for the underlying Gaussian processes of each movie, grouped by category. Horizontal and vertical components were averaged before normalizing (equivalently, each component was weighted by its variance). **E**. As in **D**, for the scale-generating Gaussian process, *Z*. **F**. The Gaussian process correlation functions in **D** averaged within categories. **G**. As in **F**, for the scale-generating Gaussian process correlation functions in **E. H**. Lag time to reach a correlation of 0.5 for the underlying velocity Gaussian processes for each movie (components were weighted by variance as in **D**). **I**. As in **H**, for the scale-generating Gaussian process. **J**. Variance explained for each movie. Variances were averaged across horizontal and vertical components before calculating *R*^2^. **K**. AIC values for for different models for each movie. Lower values indicate better model fit.

Finally, we ask whether the full ARGSM model is necessary to carry out scale normalization in practice for our trajectory data. Our model fitting provides an estimate of the scale at each time point, which we use to normalize the raw data. To quantify normalization performance, we calculate the kurtosis, or fourth standardized moment, which measures how heavy-tailed a distribution is. The standard reference is a Gaussian random variable, which has a kurtosis of 3. In Figure 8 we compare the kurtosis of the velocity before and after dividing by a point estimate of the scale under three models of increasing complexity. If normalization is successful, the distribution of the resulting normalized velocity should be approximately Gaussian. Under the time-independent model the normalized velocity consistently has kurtosis less than 3, indicating that the scale tends to be over-estimated (Figure 8A). In contrast, for a model with correlated velocity and constant scale for each trajectory, the kurtosis is consistently larger than 3, indicating that the scale tends to be underestimated (Figure 8B). Only the full model, with correlated velocity and a dynamic, correlated scale variable yields a kurtosis around 3 for each movie, even with highly kurtotic data (Figure 8C).

**FIG. 8:**
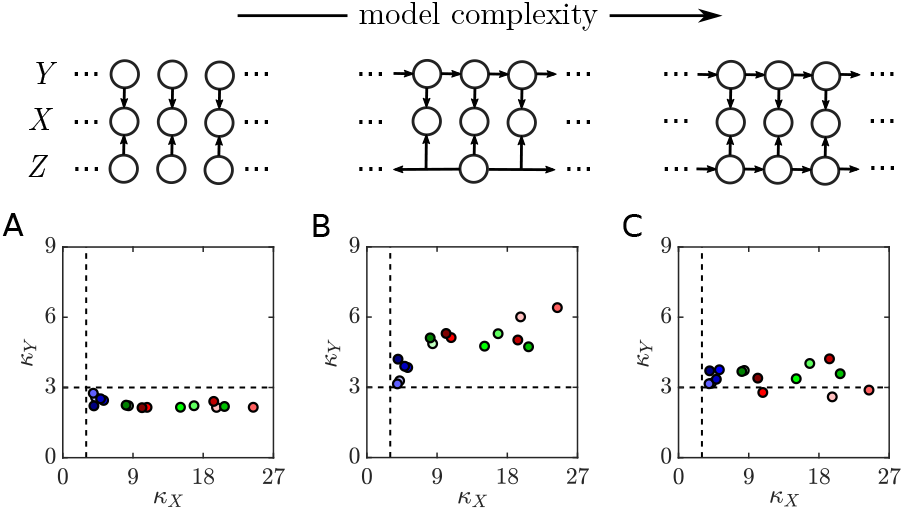
A dynamic scale-mixture model is necessary for effective normalization. **A**. Kurtosis of the velocity before and after dividing by a point estimate of the scale (*bottom*) under the time-independent model (*top*). Kurtosis was computed by pooling the two components after normalizing by each standard deviation, so that differences in the variance across components do not contribute additional kurtosis. A Gaussian distribution has a kurtosis of 3 (dashed lines). **B**. As in **A**, but for a model with autocorrelated Gaussian processes and a constant scale for each trajectory. **C**. As in **A**, but for the fully dynamic model.

This exercise of using the ARGSM model to estimate the scale at each time point, then dividing the velocity by this scale, serves as a proxy for what the nervous system can achieve through adaptation mechanisms. An important caveat is that the model has access to the full trajectory, while the nervous system must operate in an online, causal setting.

### Implications for prediction

Prediction is an important problem both for compression via predictive coding and for overcoming sensory and motor delays during behavior. Prediction is built into the ARGSM framework since the regression coefficients of the AR models are optimal for predicting the next time step of *Y*_*t*_ and *Z*_*t*_ given their past values. Let 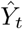 and 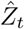 denote the predicted values:

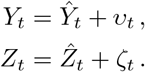

The variance explained by a prediction 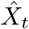 is given by

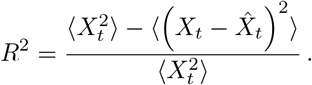

For the Gaussian process *Y*_*t*_ this simplifies to

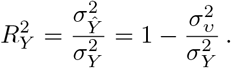

Assuming knowledge of the histories of both *X*_*t*_ and *Z*_*t*_, the prediction for *X*_*t*_ is

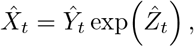

The associated variance explained is

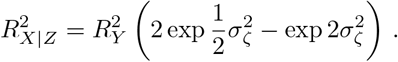

This is an upper bound on the performance of any predictor with access only to *X*_*t*_.

Notably, 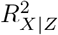 is independent of 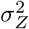 : the variance ex-plained under the ARGSM model for *X* is equal to the variance explained for *Y*, multiplied by a function of the variance of the innovation noise for *Z* that slowly decreases from one to zero. Since the innovation noise is small for the estimated models, we expect it to have little effect. In Figure 7J we compare the variance explained by applying naive autoregression to *X*_*t*_, 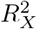, to 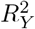 and 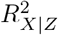 using the estimated model parameters. The variance explained is close to one for all movies except three depicting insects. Values of 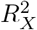 tend to be only slightly smaller than 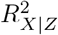. We conclude that heavy-tailed statistics have little effect on the predictability of natural motion, although scale estimation is necessary for estimating the variance associated with the prediction, that is, the variance of the posterior distribution 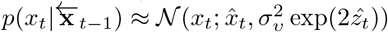.

Long correlation times and high values of *R*^2^ indicate that the velocity time series of natural motion are highly predictable. One way to make use of this predictability is through predictive coding, in which only prediction er-rors (with variance 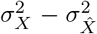, as opposed to 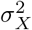 for the original signal) are sent through a channel. However, this may be a challenge for the visual system, since motion in encoded in spatial arrays of neurons rather than individual channels. A second use is actually carrying out the prediction to compensate for delays in perception or to drive motor output. Note that since the position *q* of a point is the integral of its velocity, the prediction of position by means of correlations in the velocity is given by 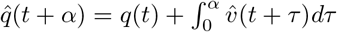, where 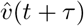 is the prediction of the velocity at time *τ* given its history up to time *t*.

## DISCUSSION

The observed pattern of heavy-tailed velocity distributions in natural movies, with a scale parameter that is shared across velocity components and fluctuates in time, is remarkably consistent across scenes and categories, despite substantial variation in the content of those scenes, velocity correlation functions, and the overall velocity variance. Together with previous results showing similar statistics in natural images and sounds [10], this suggest that scale-mixing is a fundamental property of natural stimuli, with deep implications for both neural coding and behavior.

In the context of object motion, scale-mixing may arise from two distinct mechanisms, as outlined in our discussion of Brownian motion (see *Supporting Information*). First, objects may appear at a variety of distances from the observer, and those distances may change over time. The velocity of a point on an object as it appears to an image forming device, like a camera or eye, is an angular velocity, which can be calculated as the tangential component of the physical velocity divided by the distance. A fluctuating distance thus scales the overall angular velocity over time: even an isolated point moving with Gaussian velocity statistics in three-dimensional space will have a heavy-tailed angular velocity distribution from the perspective of the observer. Second, the scale of the driving forces (either internal or external) may fluctuate over time. In our scenes, this corresponds to changes in the behavioral states of animals or to the turbulent nature of water and the wind driving plant motion. Since heavy-tailed distributions and scale fluctuations are observed in scenes with very little variance in depth, such as `bees8-full`, we emphasize that this mechanism is also at play in natural scenes.

Regardless of the source of scale-mixing, strategies for encoding and behaviorally compensating for it should be similar. On the encoding side, the presence of local scale variables suggests that sensory systems should adapt their response properties to local statistics in order to maximize information transmission. Given a fixed distribution of external input values, the optimal neural response function is the one that produces a uniform distribution over the neuron’s dynamic range [2]. The logarithmic speed tuning observed in MT [63] is consistent with this kind of static efficient coding. Here, we demonstrate that the scale of the distribution changes over time, so the gain of the response function should also change to match it [64–66]. Such adaptation or gain control is observed throughout the nervous system (see [67] for a recent review), including in systems relevant to object motion encoding[42–44, 68]. This adaptation could be the result of subcellular mechanisms, such as the molecular kinematics of synaptic vesicle release [69], or nonlinear circuit-level mechanisms [70, 71]. By measuring the timescale on which the scale variable fluctuates in natural movies scenes, we have determined the timescale on which adaptation mechanisms in the brain should operate. Although the range is considerable, for most movies the time to 0.5 correlation for the scale generator is less than one second (Figure 7I). Future experiments could be targeted at probing adaptation timescales in the retina and cortex of various model organisms that occupy different environments. Our prediction is that these adaptation variables will tightly match the motion statistics in the organism’s ecological niche.

Beyond single-cell adaptation, our results are also relevant to a population-level adaptation mechanisms known as divisive normalization [72, 73], in which neighboring neurons in a population are mutually inhibitory in a divisive fashion. In many systems, motion is represented by a local population of neurons, each tuned to a narrow band of directions. Our results show that the fluctuating scale is shared between horizontal and vertical velocity components, and, hence, adaptation should ideally be distributed throughout the local population. Divisive normalization is a prime candidate for the implementation of this population-level adaptation, as has been suggested for GSM models of filter responses [10, 74– 78]. Most models of divisive normalization only capture steady-state responses to static or constant velocity stimuli, although some work has been done to describe the dynamics of divisive normalization during change detection and decision making [79, 80]. Again, these dynamics should be tuned to the timescale of the scale fluctuations measured here.

These data suggest a previously unexplored challenge for adaptation mechanisms in the context of object motion: an object may travel an appreciable distance before local mechanisms have a chance to take effect. A solution is to pool from a larger neighborhood, or, more intriguingly, for a local population to receive an adaptation signal selectively from those neurons in nearby populations whose preferred directions point to it. To our knowledge, these hypotheses have not yet been explored, either the-oretically or experimentally.

In terms of behavior, our results help refine our understanding of the object tracking problems animals must solve in natural environments, which are crucial to survival. A commonly invoked framework for tracking is sequential Bayesian inference under a state-space model [45]. In this framework, the brain has a probabilistic representation of the state of the object (that is, a probability distribution over its position and velocity). An internal model of object motion is used to evolve this distribution forward in time, and this prediction is combined with incoming measurements to update the estimated state distribution. Under Gaussian assumptions this yields the famous Kalman filter solution [81]. Our work has two important implications for the state-space model framework of object tracking. First, the velocity distributions we observe are typically non-Gaussian, so the Kalman filter solution is not strictly applicable. While heavy tails have little impact on prediction, they have a large effect on the uncertainty of the posterior estimate. Second, state-space models usually model the velocity as either an AR(1) or (discrete) diffusion process (i.e., a nonstationary AR(1) process with coefficient equal to one). The AR models we fit for the underlying Gaussian components generally have more than one large coefficient. The ARGSM model could naturally serve as a predictive state-space model that incorporates these empirical observations by including the recent history of the velocity and scale in the state description (note that the scale does not have a corresponding direct measurement, but it can be estimated the incoming velocity measurements). Flexible Bayesian methods like the particle filter [82] can be used to implement such a model. The merging of the sort of adaptation mechanisms described above with neuromorphic particle filtering [83] is an intriguing avenue for future research.

Motion estimation itself can be framed as a Bayesian inference problem, and the tracking algorithm we use corresponds to a Gaussian prior [84]. The ARGSM model could thus serve as a better prior, motivating new motion estimation algorithms based on natural scene statistics. Speed perception in humans and animals can also be viewed through the lens of Bayesian inference, and experimental results are consistent with a heavy-tailed prior, specifically, a power law [85, 86]. The GSM model yields a heavy-tailed distribution for speed compared to the Rayleigh distribution expected under Gaussian assumptions, but it is not a true power law. Since power laws are an idealization and are always subject to some cutoff, the GSM model may be considered a more realistic (if less tractable) alternative. The correlated scale fluctuations also suggests that optimal Bayesian inference should be history-dependent, which could be assessed psychophysically using, e.g., a trial structure that is correlated in time.

Finally, the significant diversity of velocity and scale correlation functions and variances across scenes has implications both for efficient coding and tracking. Namely, an encoder or tracker which is optimized for the statistics of one scene will be suboptimal for others. Indeed, there is a general trade-off in adaptation to global versus local statistics [87, 88]. The original efficient coding work posited adaptation on evolutionary timescales to natural scene statistics. Here, we emphasize the subsecond timescale of scale fluctuations in natural motion. Neural systems should also have the flexibility to adapt on intermediate timescales to changes in the environment or behavioral context [89].

## MATERIALS AND METHODS

### Point tracking

We compute short trajectories using the PointTracker function in Matlab’s Computer Vision toolbox. The function employs a Kanade-Lucas-Tomasi [38, 39] feature tracking algorithm, which uses multi-scale image registration under a translational motion model to track individual points from frame to frame. Briefly, given an image patch *I*(*x, y, t*) centered on some seeded initial position, the algorithm finds the displacement (Δ*x*, Δ*y*) that minimizes the squared error,

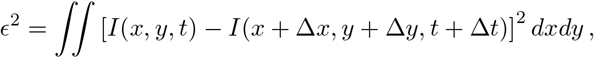

and updates the seed position on the next frame. Our strategy is to collect as many high-quality, short trajectories (64 frames) as possible from each movie, then subsample these down to a reasonable number of trajectories (8,192) for statistical analysis. Initial points are seeded using the `detectMinEigenFeatures` function, which detects image features that can be tracked well under the motion model [90]. From the initial seeds, we run the tracking algorithm forward and backward for 32 frames each, rather than running it in one direction for 64 frames. This increases the chances of capturing shortlived trajectories bounded by occlusion or image boundaries. Points are seeded on each frame, so the resulting set of trajectories is highly overlapping in time. On most movies we employ the built-in forward-backward error checking method [91], with a threshold of 0.25 pixels, to automatically detect tracking errors. The exceptions are three movies depicting water (`water3, water5`, and `water6`) where the small threshold leads to rejecting most trajectories, so we use a threshold of 8 pixels. In these cases there are not well-defined objects, so relaxing this strict criterion is justified. The algorithm uses a multi-resolution pyramid and computes gradients within a neighborhood at each level. We use the default values of 3 pyramid levels and a neighborhood size of 31 by 31 pixels for all movies except the 3 water movies, where we find we can decrease the amount of erroneously large jumps in trajectories by increasing the neighborhood size to 129 by 129 pixels and using only 1 pyramid level (at a cost of greater computation time).

This method automatically tracks the stationary background points, which may be erroneously”picked up” by a moving object as it traverses that location. To ensure that the trajectories we analyze are full of motion, we define a s peed (velocity m agnitude) t hreshold o f 0.1 pix/frame, and discard trajectories in which 16 or more time steps are below this threshold.

The velocity time series is simply the first difference of the point positions along each trajectory. Within each ensemble, we subtract the ensemble mean from each velocity component (this is typically very close to zero, except for some water movies with a persistent flow). We then slightly rotate the horizontal and vertical velocity components to remove small correlations between them (these arise if, for example, objects tended to move along a slight diagonal relative to the camera’s sensor). All visualizations and calculations are carried out after these minor preprocessing steps. Note that we do not subtract the average velocity within each trajectory, as this introduces an artificial anticorrelation at long lags.

### Gaussian scale-mixture models

The basic one-dimensional Gaussian scale-mixture model is described in the main text. Note that for some choices of the distribution for *S*, the distribution for *X* has a closed-form solution. For example, the well-known Student’s *t*-distribution is formed when *S* follows an inverse *χ*-distribution, and the Laplace distribution is formed when *S* follows a Rayleigh distribution. In this work, we assume *S* follows a log-normal distribution, which does not yield a closed-form distribution for *X*. This choice makes modeling correlations straightforward, as will be made clear below. In practice, the lack of a closed-form *p*(*x*) is not a drawback, since we do not need to normalize the posterior distribution, *p*(*s*|*x*) ∝ *p*(*x*|*s*)*p*(*s*), in order to sample from it.

When considering multiple variables, a shared scale variable introduces a nonlinear form of dependence between them. Suppose *X*_1_ = *Y*_1_*S* and *X*_2_ = *Y*_2_*S*. If *Y*_1_ and *Y*_2_ are uncorrelated, then *X*_1_ and *X*_2_ are conditionally independent given *S*:

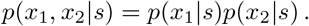

However, *X*_1_ and *X*_2_ are not, in general, independent:

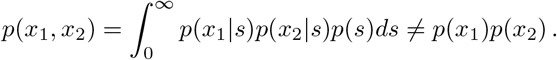

This nonlinear dependence manifests itself in the elliptic level sets of *p*(*x*_1_, *x*_2_), in contrast to the diamond-shaped level sets of *p*(*x*_1_)*p*(*x*_2_). Note that this nonlinear dependence can coincide with the usual linear dependence if *Y*_1_ and *Y*_2_ are correlated, and that a weaker form of nonlinear dependence may be present if *X*_1_ = *Y*_1_*S*_1_ and *X*_2_ = *Y*_2_*S*_2_, where *S*_1_ and *S*_2_ are not independent.

### Autoregressive models

Autoregressive models [92, 93] are a well-established and flexible way to capture correlations in time series data by supposing a linear relationship between the current value of a random variable with its previous values. Given a time series, {*X*_1_, …, *X*_*T*_ }, the *k*th order autoregressive, or AR(*k*), model is given by

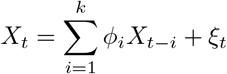

where {*ϕ*_1_, …, *ϕ*_*k*_} are regression coefficients an d *ξ*_*t*_ is Gaussian innovation noise with variance *σ*^2^.

The order *k* is typically chosen by cross-validation to avoid over-fitting. This makes sense from the standpoint of finding a m odel t hat g eneralizes w ell t o n ew data. However, our primary aim here is simply to measure the autocovariances of the hidden variables, since their timescales are relevant to prediction and adaptation in the nervous system. For this reason, we choose *k* to be as high as possible: *k* = 31 time steps, since *k* must be less than *T/*2.

Typically, the model parameters are fit by standard linear regression (after organizing the data appropriately) [94]. However, this method gives maximum likelihood estimates only if the initial *k* time steps are considered fixed. I f t he i nitial d ata a re a ssumed t o b e d rawn from the stationary distribution defined b y t he parameters, the problem becomes nonlinear. The EM algorithm described below requires parameter estimates to be maximum likelihood, and since we would like the initial *k* time steps (where *k* is large) to be modeled by the stationary distribution, we must pursue this more difficult course. We calculate the maximum likelihood estimates numerically, following [95]. See *Supporting Information* for a full description of this method.

### The ARGSM model

The dynamic scale-mixture model generalizes the two-dimensional, shared scale variable GSM model described above to time series, assuming the underlying Gaussian random variables, *Y*_1_ and *Y*_2_, and the generator, *Z*, of the scale variable are all AR(*k*) processes. Specifically, let *X*_1,*t*_ = *Y*_1,*t*_*S*_*t*_ and *X*_2,*t*_ = *Y*_2,*t*_*S*_*t*_. Written as *T* -dimensional vectors, we have **X**_1_ = **Y**_1_ ⊙ **S** and **X**_2_ = **Y**_2_ ⊙ **S**, where ⊙ is element-wise multiplication.

The AR process assumptions imply

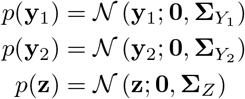

where the covariance matrices are determined by the parameters of independent AR(*k*) models as described above. **S** is related to **Z** by element-wise application of the exponential function, **S** = exp(**Z**). When **Z** is known, we have

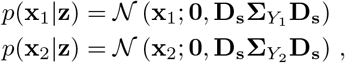

where **D**_**s**_ is the matrix with the elements of **s** along the diagonal and zeros elsewhere.

### EM and stochastic approximation

The classic Expectation-Maximization, or EM, algorithm is a useful tool for finding (local) maximum likelihood estimates of parameters of hidden variable models [55]. Let 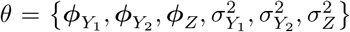 be the collec-tion of parameters of the ARGSM model, where each *ϕ* is the vector of AR coefficients and each *σ*^2^ is the innovation variance for each variable. The observed data, 𝒟 = {**x**_1,*n*_, **x**_2,*n*_}, 1 ≤ *n* ≤ *N*, are the *N* pairs of *T* - dimensional vectors corresponding here to the horizontal and vertical velocity along each trajectory. The hidden variables, ℋ = {**z**_*n*_}, 1 ≤ *n* ≤ *N*, are the Gaussian generators of the time-varying scale associated with each trajectory. The likelihood,

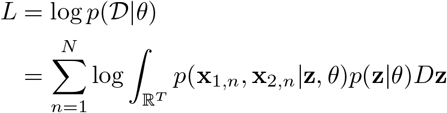

is intractable to maximize due to the high-dimensional integral. The EM algorithm finds a local maximum iteratively. Starting with an initial guess for the parameters, *θ*_0_, at each step one computes the expectation with respect to the probability distribution of the hidden variables given the data and the current parameter estimate *θ*_*t*_ of the complete data log-likelihood,

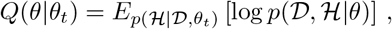

then updates the parameters to maximize this function,

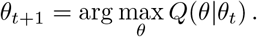

In our setting, we have

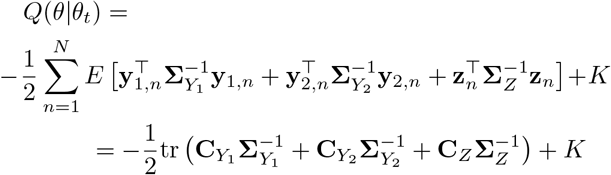

where

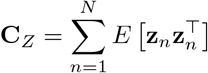

and similarly for 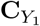 and 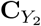. Note that in this context, the **y** variables are merely shorthand for **y**_1,*n*_ = **x**_1,*n*_ ⊘ **s**_*n*_ and **y**_2,*n*_ = **x**_2,*n*_ ⊘ **s**_*n*_, where ⊘ is element-wise division. Given the **C** matrices, the corresponding **R** matrices defined above are easily computed, from which the maximum likelihood estimates of the AR parameters can be calculated.

Unfortunately, the expectation values in the **C** matrices are also intractable, but they can be approximated through sampling methods. A variant of the EM algorithm, called stochastic approximation EM, was developed to address this problem [96]. Given a sample from the distribution *p*(ℋ| 𝒟, *θ*_*t*_), one calculates the sam-ple matrices **Ĉ**, then updates the stochastic approximations as

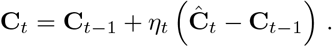

The sequence of parameters *η*_*t*_ is given by

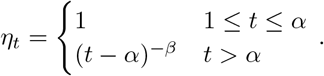

We choose *α* = 2500 or 5000, so that the algorithm runs in a fully stochastic mode until the parameter estimates are nearly stationary, and *β* = 1, so that after this initial period, the algorithm converges by simply taking a running average of the samples of the **Ĉ** matrices. Importantly, the samples do not need to be independent across iterations for the algorithm to converge [97]. This means that, when performing the Gibbs sampling described below, we only need to update each hidden variable element once for each iteration, rather than updating many times and throwing out samples to achieve independence. Since each M-step (the AR model MLE algorithm described above) is much faster than each E-step (calculating the **C** matrices through sampling), this results in a more sample-efficient algorithm [98].

We also estimate the expectation of the hidden variables {**z**_*n*_} in an identical fashion. This is equivalent to a Bayesian point estimate where the estimated parameters form a forward model and prior. These estimates are then used to remove the scale from the velocity, in order to examine the kurtosis under different model assumptions (Figure 4).

The EM algorithm, and its stochastic approximation variant, converges to a local maximum of the likelihood function that depends on the initial conditions. We find that, in practice, it is important to introduce the scale variable gradually to the model. We initialize the model with AR parameters fit to the raw data for the *Y*_1_ and *Y*_2_ components, and let *Z* be uncorrelated with very small variance (regression coefficients *ϕ*_*Z*_ = 0 and innovation variance 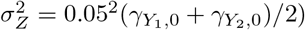.

### Sampling methods

We use a combination of Gibbs and rejection sampling to sample from the posterior of the hidden variables given the data and the current parameter estimates [99]. In Gibbs sampling, an initial vector **z** is used to generate a new sample by sampling each element individually, conditioned on the remaining elements. Since the conditional distribution is intractable, we use rejection sampling, which allows us to sample from an arbitrary, unnormalized distribution by sampling from a proposal distribution (in this case a Gaussian with parameters chosen to envelope the conditional distribution) and rejecting some draws in order to shape it into the target distribution. See *Supporting Information* for a detailed description of the sampling algorithm.

## ACKNOWLEDGEMENTS

This work was supported by the National Science Foundation through the Physics Frontier Center for Living Systems (PHY-2317138), the Center for the Physics of Biological Function (PHY-1734030), and a CAREER award to SEP (IIS-1652617); by the NSF-Simons National Institute for Theory and Mathematics in Biology, awards DMS-2235451 (NSF) and MP-TMPS-00005320 (Simons Foundation); and by the National Institutes of Health BRAIN Initiative (R01EB026943). We thank Siwei Wang and Benjamin Hoshal for useful comments on the manuscript.

