## Supporting Information for "A dynamic scale-mixture model of motion in natural scenes"

### I. A MOTIVATING EXAMPLE: BROWNIAN MOTION

Here we introduce a canonical example of natural motion to guide our intuition about departures from the simplest case: Brownian motion. Brown discovered the motion that bears his name by observing small particles released by a grain of pollen floating in water under a microscope [1]—a highly controlled setting, but similar in spirit to our own. Each particle is subjected to tiny molecular forces from individual water molecules colliding with it at random. As a result, it moves in a random fashion across the surface of the water, limited only by the boundary of the dish. Its motion is not completely unpredictable, however; since the particle has some mass, albeit small, it has some inertia, or tendency to continue moving with the same velocity. Statistically, this means the velocity is correlated in time.

Based on this observation, Einstein derived his famous diffusion equation, describing how a density of diffusing particles changes over time, by considering the case of particles with infinitesimal mass [2]. For describing the motion of an individual, massive particle, it is more useful to look to Langevin's description, an early application of stochastic differential equations [5]. Using Newton's law and assuming the force can be divided into the sum of a drag force, proportional to the velocity by a factor of $-\gamma$, and a randomly fluctuating force due to the collision of water molecules, $F(t)$, we have the following differential equation for the velocity, $V(t)$, of a particle with mass $m$:

$$m\frac{d}{dt}V(t) = -\gamma V(t) + F(t)\,.$$

When $F(t)$ is an uncorrelated Gaussian process, $V(t)$ is an Ornstein-Uhlenbeck process, which has an exponential correlation function (see [3, 4] for a pedagogical review). The variance of the velocity, $\sigma^2$, is related to the temperature, $T$, by $\sigma^2 = kT/m$, where $k$ is Boltzmann's constant.

Numerically and experimentally, we must always discretize time. The discrete-time approximation to the Ornstein-Uhlenbeck process with time-step $\Delta t$ is given by the difference equation

$$V_{t+\Delta t} = \phi V_t + \xi_t\,,$$
$$p(\xi_t) = \mathcal{N}(\xi_t; 0, \sigma_\xi^2)\,,$$

which is a first-order autoregressive, or AR(1), process [8, 9]. Like the Ornstein-Uhlenbeck process, it has an exponential autocovariance function, given by

$$E\left[V_t V_{t+k\Delta t}\right] = \sigma^2 \exp\left(-\frac{k\Delta t}{\tau}\right) ,$$

with variance

$$\sigma^2 = \frac{\sigma_\xi^2}{1-\phi^2}$$

and time constant

$$\tau = -\frac{\Delta t}{\ln \phi} .$$

Let us consider how to incorporate a fluctuating variance in the AR(1) process. There are two possibilities: the variance of the driving noise $\xi_t$ could fluctuate, as in Brownian motion with a fluctuating temperature. Alternatively, the entire process could be scaled by a fluctuating positive variable; in the Brownian motion experiment, this corresponds to changes in the magnification level of the microscope. This also corresponds to changes in inverse distance if the velocity is an angular velocity relative to an observer. The former, additive model corresponds to

$$V^+_{t+\Delta t} = \phi V^+_t + S_t \xi_t .$$

The latter, multiplicative model corresponds to

$$V^\times_t = S_t V_t ,$$

where $S_t$ is the fluctuating scale. Rewriting

$$V^\times_{t+\Delta t} = \frac{S_{t+\Delta t}}{S_t} \phi V^\times_t + S_{t+\Delta t} \xi_t ,$$

we see that $V^+_t$ and $V^\times_t$ are identical when $S_t$ is constant in time and varies only across the ensemble. However, they are not equivalent when $S_t$ changes over time. In the main text, we adopt the multiplicative model, since it factors into a product of two independent stochastic processes (one for normalized velocity and one for scale), which simplifies the inference problem.

### II. TRACKING ALGORITHM VALIDATION

We validated the tracking algorithm by generating a synthetic movie, computing trajectories, and comparing them to ground truth. The synthetic movie consists of a single frame from `bees8-full` with a grid of 16 disks (of radius 48 pixels) copied from their initial positions, overlaid

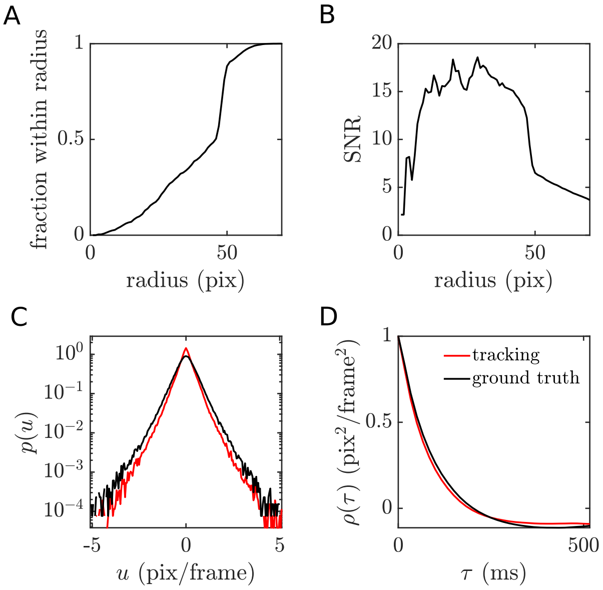


FIG. 1: **Tracking performance.** **A**. Fraction of trajectories within a given radius of the center of the moving disks. **B**. Tracking signal-to-noise ratio for the subset of trajectories within a given radius. **C**. Horizontal velocity histogram for the tracking data and ground truth. **D**. Horizontal velocity autocorrelations for the tracking data and ground truth.

on the original image, and animated according to an ARGSM process. The underlying Gaussian velocity time series were generated by simulating critically damped harmonic oscillators (independently for each component) with velocity standard deviation $\sigma_{Y_1} = \sigma_{Y_2} = 0.5$ and time constant $\tau = 100$ ms. The scale time series were generated by simulating an AR(1) process with standard deviation $\sigma_Z = 0.5$ and coefficient $\phi_Z = 0.75$. We multiplied each pair of velocity components by the scale and calculated the cumulative sum to determine the position of the disks on each frame relative to their grid positions. The disks occasionally occlude one another but are largely isolated; we expect that adding more disks and occlusions decreases tracking performance accordingly.

We apply the tracking algorith to generate 8,192 trajectories of length 65. After applying the tracking algorithm, we associate each tracking trajectory with a given ground truth trajectory if its initial position is within a given radius of the center of the corresponding disk; trajectories which are not associated with any ground truth are discarded. We calculate the mean squared error of the velocity along these trajectories relative to the ground truth velocity at the corresponding times. Looking at the fraction that remains (Figure 1A), we find that many trajectories are seeded at the perimeter of the disk, since there tends to be a distinct edge there. The presence of trajectories

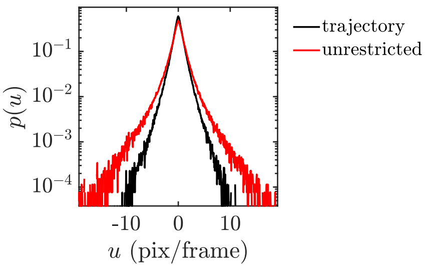


FIG. 2: **Velocity histograms.** Histograms for horizontal velocity with and without restriction to trajectories.

with initial positions outside of any disk indicates that the algorithm detects some spurious motion in the vicinity of the actual motion. We calculate the mean squared error of the velocity along these trajectories relative to the ground truth velocity at the corresponding times. The signal-to-noise ratio (SNR) (Figure 1B), or ratio of the variance of the ground truth velocity to the mean squared error, drops off quickly when the radius is large enough to include the spurious trajectories. However, the statistics of the ensemble are very close to the ground truth when all of the tracking trajectories are included. The distribution of a single velocity component (Figure 1C) captures the overall shape and tails of the ground truth distribution but is more sharply peaked. The autocorrelation (Figure 1D) is underestimated only slightly.

### III. EFFECT OF RESTRICTING VELOCITY TO TRAJECTORIES

Since tracking is a difficult problem that involves associating velocity values in time subject to our criteria for trajectory quality, the set of velocity values along trajectories is only a subset of all velocity values in a given scene. To understand the difference, we apply the same algorithm with the trajectory length set to two, so there is only single velocity per trajectory, with no forward-backward error thresholding. For `bees8-full`, we find a substantially broader distribution for the unrestricted velocity (see Figure 2). The sample standard deviation and kurtosis for the trajectory-restricted velocity are 1.2 pix$^2$ and 10.2, respectively, compared to 1.9 pix$^2$ and 24.5 for the unrestricted velocity. We thus expect the standard deviation and kurtosis values reported here to be underestimates compared to theoretically perfect tracking.

### IV. MAXIMUM LIKELIHOOD ESTIMATION FOR AUTOREGRESSIVE MODELS

Following [6], we consider the time series as a $T$-dimensional vector, $\mathbf{X} = [X_1 \dots X_T]^\top$. Then

$$p(\mathbf{x}) = \mathcal{N}(\mathbf{x}; \mathbf{0}, \mathbf{\Sigma}) \,,$$

where

$$\mathbf{\Sigma} = E[\mathbf{X}\mathbf{X}^\top] = \text{Toeplitz}(\gamma_0, \dots, \gamma_{T-1}) \,.$$

The autocovariance series $\{\gamma_0, \dots, \gamma_{T-1}\}$ is related to the regression coefficients and innovation variance by

$$\gamma_0 = \sum_{i=1}^{k} \phi_i \gamma_i + \sigma^2$$
$$\gamma_j = \sum_{i=1}^{k} \phi_i \gamma_{i-j} \qquad j \geq 1 \,.$$

Given a collection of $N$ samples of length $T$ from the time series, we calculate the matrix $\mathbf{R}$ defined by

$$R_{ij} = R_{ji} = \frac{1}{N} \sum_{n=1}^{N} \sum_{t=i+1}^{T-j} x_{n,t} x_{n,t+i-j} \qquad 0 \leq i \leq j \leq k \,,$$

where the index $n$ ranges over samples and $t$ ranges over time steps. Then the maximum likelihood estimate of the parameters satisfies a nonlinear system of $2(k+1)$ equations, consisting of the $k+1$ autocovariance relations above, up to $\gamma_k$, together with the following:

$$R_{00} = \sum_{i=1}^{k} \phi_i \left( R_{0i} + i\gamma_i \right) + T\sigma^2$$
$$R_{j0} = \sum_{i=1}^{k} \phi_i \left( R_{ij} + i\gamma_{i-j} \right) \qquad 1 \leq j \leq k \,.$$

We solve this system numerically using Matlab's `fsolve`.

While the covariance matrix is easily calculated from the AR model parameters, it is not well-appreciated that the inverse covariance matrix can be calculated exactly from the parameters as well [7]. This is useful because it is the inverse that is needed for the sampling methods described below, and inverting large, nearly singular matrices is numerically unstable. The inverse of $\mathbf{\Sigma}$ is given by

$$\mathbf{\Sigma}^{-1} = \frac{1}{\sigma^2} \left( \mathbf{I} + \sum_{i=1}^{k} \phi_i^2 \mathbf{E}_i - \sum_{i=1}^{k} \phi_i \mathbf{F}_i + \sum_{i=1}^{k-1} \sum_{j=1}^{p-i} \phi_i \phi_{i+j} \mathbf{G}_{j,i+j} \right) \,,$$

where $\mathbf{I}$ is the identity matrix, $\mathbf{E}_i$ is the identity matrix with the first and last $i$ diagonal elements set to zero, $\mathbf{F}_i$ is a matrix with ones along the $i$th upper and lower diagonals and zeros elsewhere, and $\mathbf{G}_{j,i+j} = \mathbf{E}_i \mathbf{F}_j \mathbf{E}_i$ (note that we have corrected a typo in [7] in the indexing of the last term of the equation).

### V. SAMPLING METHODS

Gibbs sampling works by starting with an initial vector $\mathbf{z}$ and updating each element $z_t$ (in a random order) conditioned on the remaining elements $\mathbf{z}_{\setminus t}$, where the notation $\setminus t$ indicates all indices in the set $\{1, \ldots, T\} \setminus t$. The posterior distribution is given by

$$
\begin{aligned}
& p(z_t|\mathbf{x}_1, \mathbf{x}_2, \mathbf{z}_{\setminus t}) \propto \\
& \quad p(x_{1,t}, x_{2,t}|\mathbf{x}_{1,\setminus t}, \mathbf{x}_{2,\setminus t}, \mathbf{z}) p(z_t|\mathbf{x}_{1,\setminus t}, \mathbf{x}_{2,\setminus t}, \mathbf{z}_{\setminus t}) \\
& \qquad = p(x_{1,t}|\mathbf{x}_{1,\setminus t}, \mathbf{z}) p(x_{2,t}|\mathbf{x}_{2,\setminus t}, \mathbf{z}) p(z_t|\mathbf{z}_{\setminus t}) \, .
\end{aligned}
$$

Each distribution in the product is a Gaussian given by

$$
\begin{aligned}
p(x_{1,t}|\mathbf{x}_{1,\setminus t}, \mathbf{z}) &= \mathcal{N}(x_{1,t}, \hat{\mu}_{Y_1} s_t, \hat{\sigma}^2_{Y_1} s_t^2) \\
p(x_{2,t}|\mathbf{x}_{2,\setminus t}, \mathbf{z}) &= \mathcal{N}(x_{2,t}, \hat{\mu}_{Y_2} s_t, \hat{\sigma}^2_{Y_2} s_t^2) \\
p(z_t|\mathbf{z}_{\setminus t}) &= \mathcal{N}(\hat{\mu}_Z, \hat{\sigma}^2_Z) \, ,
\end{aligned}
$$

where

$$
\begin{aligned}
\hat{\sigma}^2_{Y_1} &= \left(\boldsymbol{\Sigma}^{-1}_{Y_1}\right)^{-1}_{tt} \\
\hat{\sigma}^2_{Y_2} &= \left(\boldsymbol{\Sigma}^{-1}_{Y_2}\right)^{-1}_{tt} \\
\hat{\sigma}^2_Z &= \left(\boldsymbol{\Sigma}^{-1}_Z\right)^{-1}_{tt} \\
\hat{\mu}_{Y_1} &= -\hat{\sigma}^2_{Y_1} \left(\boldsymbol{\Sigma}^{-1}_{Y_1}\right)_{t,\setminus t} \mathbf{y}_{1,\setminus t} \\
\hat{\mu}_{Y_2} &= -\hat{\sigma}^2_{Y_2} \left(\boldsymbol{\Sigma}^{-1}_{Y_2}\right)_{t,\setminus t} \mathbf{y}_{2,\setminus t} \\
\hat{\mu}_Z &= -\hat{\sigma}^2_Z \left(\boldsymbol{\Sigma}^{-1}_Z\right)_{t,\setminus t} \mathbf{z}_{\setminus t} \, .
\end{aligned}
$$

Rejection sampling is used to draw a sample of each individual $z_t$. In this framework, a proposal distribution that is easy to sample from, $q(z)$, is chosen as an envelope (after an appropriate scaling) of a target distribution, $\hat{p}(z)$, from which we would like to sample, that is, $\hat{p}(z) \leq Mq(z)$ for some positive scaling factor $M$. The proposal distribution is sampled from, followed by sampling another variable $u$ from a uniform distribution. If $u \leq Mq(z)/\hat{p}(z)$, the sample is accepted; otherwise, it is rejected and the sampling is repeated. The method produces samples from the target distribution exactly, even if it is unnormalized, but requires the envelope to be tight to avoid rejecting too many samples. Here, the target distribution is the the unnormalized posterior (dropping time indices for clarity),

$$\hat{p}(z) = \mathcal{N}(x_1; \hat{\mu}_{Y_1} s, \hat{\sigma}^2_{Y_1} s^2)\mathcal{N}(x_1; \hat{\mu}_{Y_2} s, \hat{\sigma}^2_{Y_2} s^2)\mathcal{N}(z; \hat{\mu}_Z, \hat{\sigma}^2_Z)\,.$$

We let the proposal distribution be a Gaussian,

$$q(z) = \mathcal{N}(z; \mu_q, \sigma_q^2)\,,$$

whose mean and variance are optimized heuristically to form a tight envelope of the target distribution. We center this Gaussian over the peak of the target distribution,

$$\mu_q = \arg\max_z \hat{p}(z)\,.$$

and compute the scale factor $M$ to match the two distributions at the peak, $M = \hat{p}(\mu_q)/q(\mu_q)$. To optimize the variance, note that we can write

$$\begin{aligned} &\log \hat{p}(z) = \\ &-\frac{1}{2}\frac{\left[x_1 \exp(-z) - \hat{\mu}_{Y_1}\right]^2}{\hat{\sigma}^2_{Y_1}} - \frac{1}{2}\frac{\left[x_2 \exp(-z) - \hat{\mu}_{Y_2}\right]^2}{\hat{\sigma}^2_{Y_2}} \\ &-\frac{1}{2}\frac{\left[z - \left(\hat{\mu}_Z - 2\hat{\sigma}^2_Z\right)\right]^2}{\hat{\sigma}^2_Z} - 2\left(\hat{\mu}_Z - \hat{\sigma}^2_Z\right) + K\,. \end{aligned}$$

The first two terms rapidly approach constants to the right and diverge negatively to the left. Thus, the right tail of $\hat{p}(z)$ behaves as $\mathcal{N}(z; \hat{\mu}_Z - 2\hat{\sigma}^2_Z, \hat{\sigma}^2_Z)$, and the left tail falls off extremely rapidly. A variance of $\sigma^2_q = \hat{\sigma}^2_Z$ is necessary and sufficient to cover the two tails when the peak of the proposal distribution is to the right of the peak of this Gaussian distribution, i.e., when $\mu_q \geq \hat{\mu}_Z - 2\hat{\sigma}^2_Z$. A smaller variance will not fully envelop the right tail regardless of the value of $\mu_q$. We require a slightly larger variance when $\mu_q < \hat{\mu}_Z - 2\hat{\sigma}^2_Z$, which we calculate by assuming $Mq(z)$ makes exactly one other point of contact with $\hat{p}(z)$. Let

$$f(z) = \log \hat{p}(z)$$

and

$$\begin{aligned} g(z) &= \log[Mq(z)] \\ &= -\frac{1}{2}\frac{(z-\mu_q)^2}{\sigma_q^2} + f(\mu_q) \end{aligned}$$

Let $z_0$ be the point of contact. Then $f(z_0) = g(z_0)$ implies

$$\sigma_q^2 = \frac{1}{2} \frac{(z_0 - \mu_q)^2}{f(\mu_q) - f(z_0)} .$$

The two curves must also be tangent at $z_0$, $f'(z_0) = g'(z_0)$. We have

$$g'(z_0) = -\frac{z_0 - \mu_q}{\sigma_q^2} = -2\frac{f(\mu_q) - f(z_0)}{z_0 - \mu_q} .$$

The tangency condition implies $z_0$ is the solution to

$$f'(z_0) + 2\frac{f(\mu_q) - f(z_0)}{z_0 - \mu_q} = 0 ,$$

which we solve numerically using Matlab's `fzero` function, and use this solution to calculate $\sigma_q^2$. Finally, if the above procedure fails due to numerical issues, we simply set $\sigma_q^2 = (1.1\hat{\sigma}_z)^2$, which is large enough to form an envelope in practice.

---


[1] Robert Brown. A brief account of microscopical observations made in the months of June, July and August, 1827, on the particles contained in the pollen of plants; and on the general existence of active molecules in organic and inorganic bodies. *The Philosophical Magazine*, 4(21):161–173, 1828.
